# Wild flies hedge their thermal preference bets in response to seasonal fluctuations

**DOI:** 10.1101/2020.09.16.300731

**Authors:** Jamilla Akhund-Zade, Denise Yoon, Alyssa Bangerter, Nikolaos Polizos, Matthew Campbell, Anna Soloshenko, Thomas Zhang, Eric Wice, Ashley Albright, Aditi Narayanan, Paul Schmidt, Julia Saltz, Julien Ayroles, Mason Klein, Alan Bergland, Benjamin de Bivort

## Abstract

Fluctuating environmental pressures can challenge organisms by repeatedly shifting the optimum phenotype. Two contrasting evolutionary strategies to cope with these fluctuations are 1) evolution of the mean phenotype to follow the optimum (adaptive tracking) or 2) diversifying phenotypes so that at least some individuals have high fitness in the current fluctuation (bet-hedging). Bet-hedging could underlie stable differences in the behavior of individuals that are present even when genotype and environment are held constant. Instead of being simply ‘noise,’ behavioral variation across individuals may reflect an evolutionary strategy of phenotype diversification. Using geographically diverse wild-derived fly strains and high-throughput assays of individual preference, we tested whether thermal preference variation in *Drosophila melanogaster* could reflect a bet-hedging strategy. We also looked for evidence that populations from different regions differentially adopt bet-hedging or adaptive-tracking strategies. Computational modeling predicted regional differences in the relative advantage of bet-hedging, and we found patterns consistent with that in regional variation in thermal preference heritability. In addition, we found that temporal patterns in mean preference support bet-hedging predictions and that there is a genetic basis for thermal preference variability. Our empirical results point to bet-hedging in thermal preference as a potentially important evolutionary strategy in wild populations.

## Introduction

Individuals differ in their behavior – these differences are primarily caused by variation in environment, age, sex, and genetics. Many behaviors, in many species, have been found to be consistent across environments and time at an individual level^1–5^. The ubiquity of such stable variability is evidence for potential ecological and evolutionary importance, and the origins of this variability at the genetic and evolutionary levels are subjects of ongoing research.

Temporal fluctuations in the environment can lead to the maintenance of interindividual differences in natural populations. At different points in time, environmental pressures will select for different optimum phenotypes^6–9^. If interindividual differences are determined by genetic polymorphisms segregating in the population, polymorphism frequencies will change due to adaptation to the new selection pressures. As a consequence, the average behaviors of individuals may also change over time^10,11^. This process is referred to as adaptive tracking^12^, and in populations using this strategy, phenotypes lag selective pressures. When fluctuations are relatively rapid, selective pressures may be reversed by the time a population adapts, which can lead to lower fitness^13–15^. In such situations, it can be advantageous to decouple genetic and phenotypic variation, thereby reducing the phenotypic mismatch when selective pressures and adaptive phenotypic responses are out of phase^15^.

Diversifying bet-hedging (bet-hedging herein) is an alternative strategy that can overcome the limitations of adaptive tracking in rapidly fluctuating environments, because it makes it likely that there will always be at least some fit individuals for any state of the environment^7,16–18^. Under this strategy, a single genotype produces multiple phenotypes as a way to mitigate risk i.e., ‘don’t put all your eggs in one basket’. This bet-hedging strategy reduces fitness variance across generations, increasing geometric mean fitness at the expense of arithmetic fitness^16,19–21^. Intuitively, this means that although in a single generation a bet-hedging population may not be optimally fit for a given environment, the stability of fitness over generations (a bet-hedger is less likely to drop to zero fitness, extinction, at any point) results in long term success. As environmental variance increases, bet-hedging becomes an evolutionarily optimal strategy that can explain the maintenance of inter-individual differences^22–26^. There is a variety of evidence for bet-hedging traits across organisms^12^, though there are few examples of bet-hedging in animals or behavioral phenotypes.

We expect that individuals from a bet-hedging genotype would exhibit stable idiosyncratic behavioral biases when reared in the same environment. Isogenic *Drosophila melanogaster*, reared in a constant lab environment, exhibit consistent, but widely varying, individual behaviors — examples include turning bias^27–29^, phototaxis^30^, thermal preference^12^, and odor preference^31^. These differences are termed intragenotypic variability, and they may reflect a bet-hedging strategy. Kain *et al*. used a model that translated observed thermal preference intragenotypic variability into simulated variability in life-history, under either a bet-hedging or adaptive-tracking strategy, in simulated populations of *Drosophila melanogaster*^32^. They found that bet-hedging is more advantageous than adaptive tracking in environments with a high variance in seasonal temperatures and a short breeding season^32^. Despite this effort, there is scarce empirical, hypothesis-driven evidence that a bet-hedging strategy explains individual variability in animal behavior.

We conducted several empirical tests of the predictions made by an updated temperature-dependent fly life-history model. These experiments tested the hypotheses that 1) thermal preference in *Drosophila melanogaster* follows a bet-hedging strategy, and 2) bet-hedging and adaptive tracking strategies vary geographically. Measuring the thermal preferences of many individual flies collected wild from multiple locations across the USA, we found that 1) patterns of mean thermal preference over time are more consistent with bet-hedging than adaptive tracking, and 2) across geographic locations there is variation in thermal preference variability (which was uncorrelated with the predicted bet-hedging advantage) as well as variation in the heritability of thermal preference (which was negatively correlated with the predicted bet-hedging advantage, as expected).

## Results

### Individual thermal preference is idiosyncratic and persistent

If individual variation in thermal preference behavior reflects an evolutionary bet-hedging strategy, individual preferences would represent phenotypes that are stable on relatively long timescales. We created a two-choice preference assay to measure individual thermal preferences of many flies in parallel (Fig. 1a). We measured an individual fly’s thermal preference as the time-weighted average of the temperatures experienced as it navigates the hot and cold sides of the assay (Fig. 1b). We compared the observed distribution of thermal preferences within a wild-type line of isogenic animals raised in a temperature-controlled incubator to the null distribution expected from flies with identical preferences. In such a null model, sampling error alone generates the dispersion in a measured distribution (null distribution was generated by bootstrap resampling of walking bouts; see Methods and Honegger & Smith, *et al*. 2019^31^). The observed distribution was significantly broader than the null distribution (Kol-mogorov-Smirnov test, *p* = 9.1×10^−9^; Fig. 1c), indicating that thermal preferences are over-dispersed, i.e., the flies exhibited individuality in thermal preference. We also observed a significant positive correlation (Fig. 1d; *r* = 0.50, *p* = 6.2×10^−5^) in thermal preference measured on consecutive days, consistent with individual measured thermal preferences representing stable phenotypes. The repeatability of thermal preference, calculated as the intraclass correlation coefficient^33^ was 0.47, which is typical of repeatabilities for behavioral traits across a variety of organisms^34^.

**Figure 1.**
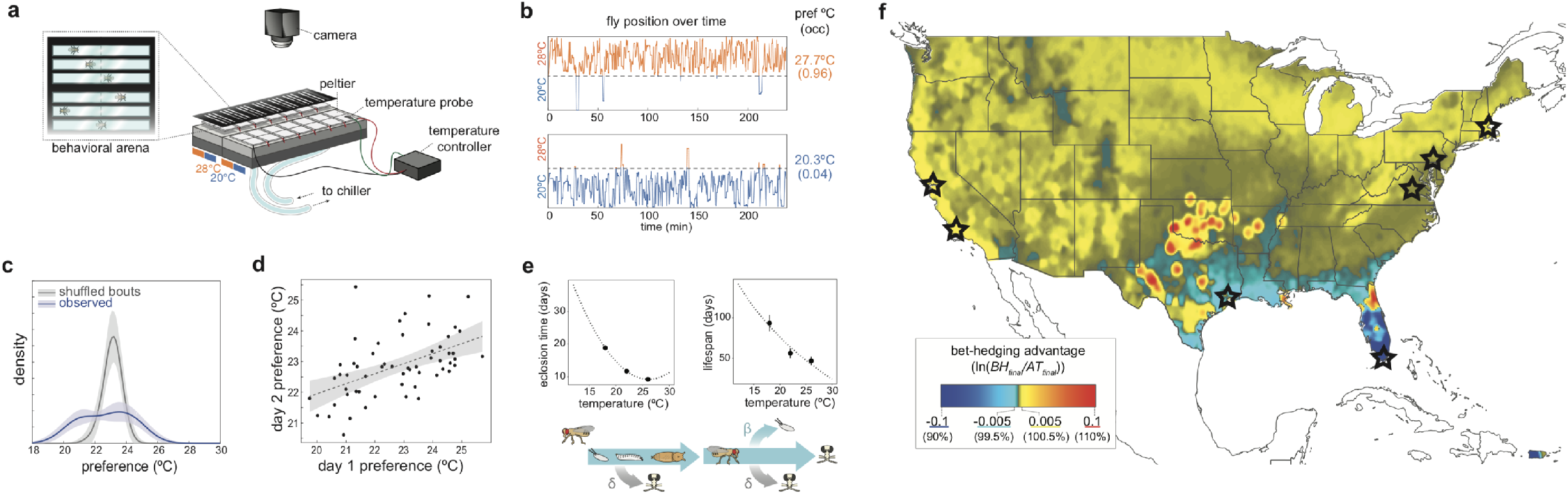
Measuring stable individual thermal preference and model predictions of the advantage of bet-hedging in thermal preference. **a)** Diagram of the two-choice assay used to measure individual thermal preference. **b)** Examples of position vs. time in the assay for extreme cold- and warm-preferring flies. Thermal preference (pref °C) is calculated as the time-averaged temperature experienced by each fly. Shown in parentheses is the fraction of time spent on the hot side. **c)** Kernel density estimates of the observed (blue) and null distribution (gray) of thermal preference for an isogenic line. Shaded areas show +/− 2 s. d. of kernel-density estimates from 100 bootstrap resamples of observed data (*n* = 57) or 100 simulated null distributions. **d)** Persistence of individual thermal preference over 24 hrs in an isogenic line (*n* = 57). Shaded area shows the 95% confidence interval (CI) of a linear fit to the data. **e) (top)** Life-history vs. temperature relationships used in the bet-hedging vs. adaptive tracking model. Error bars show show +/− 2 s. e. m. **(bottom)** Temperature dependent life-history model used to simulate fly populations (adapted with permission from Kain *et al*.^32^): β, birth rate; δ, death rate. “Fly skull and crossbones” icons indicate death. **f)** Map of bet-hedging advantage across the continental USA and Puerto Rico calculated using a Gaussian convolution of the predicted bet-hedging advantage at 7112 weather stations. Field sites for experiments below are overlaid in black stars.

### Life-history modeling predicts that the adaptive value of bet-hedging varies geographically

To generate specific predictions about the thermal preference behavior of wild flies implementing a bet-hedging strategy, we turned to a temperature-dependent life-history model of fly development and mortality^32^ (Fig. 1e). This model estimates the dynamics of populations implementing pure bet-hedging or adaptive-tracking strategies, under specific temperature profiles. At its core, the model subjects flies to a tradeoff between faster development but shorter adult lifespan (and therefore lower lifetime fecundity), and slower development with an incurred elevated risk of dying before sexual maturity. Since temperature preference affects both development time and adult lifespan^35–37^, individual flies’ thermal behavior determines these life-history traits. We improved the model in two ways: 1) we updated the temperature-dependent life-history equations with fits to new data collected from isofemale lines established using wild-caught females (Fig. 1e) and 2) we implemented more realistic rules to convert individual thermal preference, in combination with the range of temperatures available on a given day, to individual thermal experience (See Methods). With these improvements, we predicted the relative advantage of bet-hedging vs. adaptive tracking (Fig. 1f) across the continental USA and Puerto Rico using the 1981-2010 climate normals (typical daily temperature profiles) from 7112 weather stations (US National Oceanic and Atmospheric Administration [NOAA]^38^). To make predictions at sites between weather stations, and take into account dispersal of flies, we computed the 2D Gaussian convolution of the station data with a standard deviation of 0.04 decimal degrees (equivalent to 3-4km depending on the latitude). We chose this value based on empirical data from a release and recapture study^39^ that found appreciable frequencies (~15%) of marked flies 3-6km from the release site.

**Figure 2.**
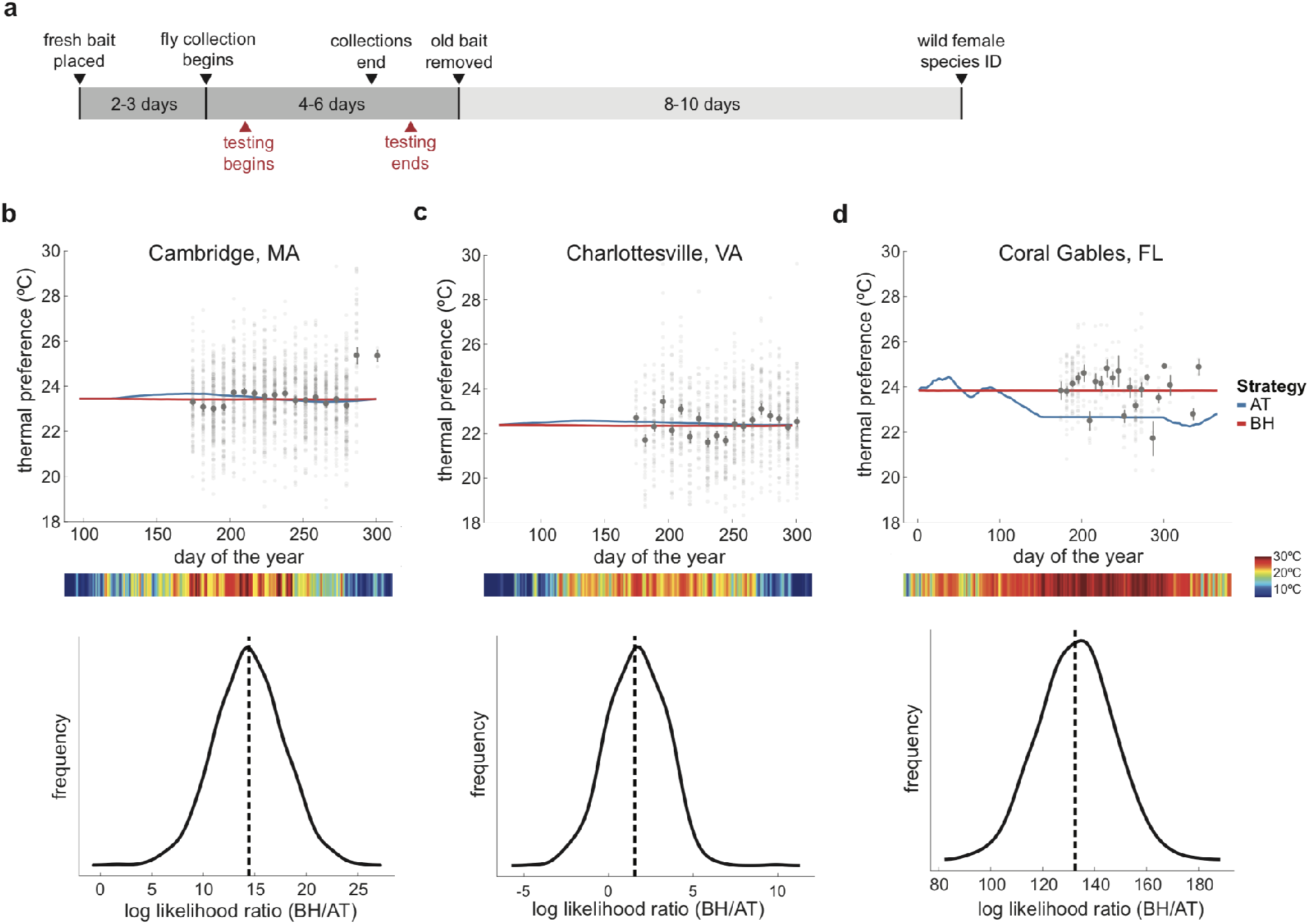
Dynamics of mean thermal preference over the course of the fly breeding season. **a)**Timeline for the seasonal sampling. For further detail see Supp. Fig 6. **b) (top)** Thermal preference of flies collected in MA. Solid lines show the predicted mean thermal preference under bet-hedging (BH, red) and adaptive tracking (AT, blue). To create these predictions, we used 2018 daily average temperatures^38^ from April to November and empirically determined location-specific behavioral means and variances. Light gray points are individual flies, with the dark gray point and error representing the mean +/− 1 s. e. m of that week’s collection. Heat map below the x-axis is the temperature of each day. **(bottom)** Log likelihood ratio (BH/AT) of bet-hedging and adaptive-tracking models given the observed mean thermal preferences. Solid line is the kernel-density estimate of the log likelihood ratio as calculated from 1000 bootstrap resamples. **c)** As in (b) for flies collected in VA at Carter Mountain Orchard. **d)** As in (b) for flies collected on campus and in residential neighborhoods next to University of Miami in FL.

Bet-hedging advantage in each region was calculated as the natural log of the final size of the simulated bet-hedging population over the final size of the simulated adaptive tracking population (*BH_adv_* = ln(*BH_pop, final_*/*AT_pop, final_*)), with each simulation run separately. Overall, the model predicted that bet-hedging advantage is generally greater at higher latitudes. In the western half of the USA, bet-hedging was predicted as advantageous even in southern latitudes. In the eastern half of the USA, we predicted that adaptive tracking is favored over bet-hedging in much of Texas, the Gulf Coast, Florida and Puerto Rico. Notably, within these regions, the model predicted heterogeneity in bet-hedging advantage, likely due to local microclimates.

### Seasonal dynamics of mean thermal preference are consistent with a bet-hedging strategy

Because bet-hedging populations are not responsive to natural selection, their mean phenotypes may be more stable over time. We hypothesized that flies from locations where bet-hedging is predicted to be advantageous would exhibit more stable mean thermal preference dynamics. To test our hypothesis, we assayed wild-caught flies weekly from late June to late October/early November in Cambridge, Massachusetts, USA (MA), Charlottesville, Virginia, USA (VA), and Coral Gables, Florida, USA (FL). Flies were captured in residential areas of MA and FL and in an orchard on the outskirts of VA; assays were performed in a laboratory environment near each collection site (Fig. 2a, Methods). We chose these locations due to their differences in predicted bet-hedging advantage.

The vast majority of bet-hedging vs adaptive-tracking predictions for the USA were within 1% annual growth advantage or disadvantage for the bet-hedging strategy. Compounded over years, this annual growth advantage can have a strong impact on the success of a population. Of the three sites where we collected seasonal dynamics data, MA is predicted to be the most bet-hedging advantageous (BH advantage = 0.0042, corresponding to an annual growth advantage of 0.42% for a bet-hedging strategy), followed by VA (BH advantage = 0.0020, annual growth advantage of 0.20% for a bet-hedging strategy), while FL is strongly favored for adaptive tracking (BH advantage = −0.54, annual growth disadvantage of 58% for a bet-hedging strategy) (Fig. 1f). To test whether the seasonal patterns in mean preference followed bet-hedging or adaptive-tracking predictions, we plugged daily temperature data from 2018 into our model to generate site-and-year-specific predicted patterns in mean thermal preference for a purely bet-hedging or a purely adaptive tracking population. We calculated a log-likelihood ratio to gauge whether the observed dynamics are more likely under a bet-hedging or an adaptive tracking strategy.

We found that in MA, the dynamics of mean thermal preference were more consistent with bet-hedging than adaptive tracking (log-likelihood ratio [LLR] = 14.4; Fig. 2b). This was consistent with our modeling predictions. In VA, the observed data were still more likely under a bet-hedging strategy (LLR = 1.56; Fig. 2c), though the relative likelihood of the bet-hedging model was less than in MA. Consistently, our model predicted VA to be bet-hedging favored, though less so than MA. In FL, the observed dynamics of mean thermal preference are much more likely under the bet-hedging than the adaptive tracking model (LLR = 133; Fig. 2d). However, our model predicts that flies in FL will exhibit adaptive tracking, including a clear selection for colder thermal preference. Interestingly, the model predicted a steep decline in total population during the hottest part of the summer, a pattern that was to some degree evident in our collections (similar seasonal population declines were observed in southern latitude populations of *D. subobscura*^37^). We observed high heterogeneity in the number of flies collected per week, with a sharp drop roughly coinciding with the bottoming out of the model (Supp. Fig. 1).

It is possible that the mean preference dynamics we observed were the result of plastic responses to ambient temperatures. We assessed the potential role of developmental temperature on mean preference by rearing flies from a single isofemale line at 18°C, 22°C, and 26°C from egg to adulthood. Rearing flies at different temperatures had a small (~0.6°C) and non-linear effect on the preference mean and standard deviation (Supp. Fig. 2), suggesting that temperature-induced plasticity in thermal preference is not the major cause of the observed dynamics.

### Variability in thermal preference is under genetic control, but is uncorrelated with predicted bet-hedging advantage

We expected that genotypes from bet-hedging populations would show higher variability than those from adaptive-tracking populations, as the latter can undergo purifying selection. Under adaptive tracking, deviations from the mean phenotype are maladaptive if a genotype is adapted to the current environment. In contrast, phenotypic diversity is an essential part of the diversification strategy of bet-hedging, and therefore would be maintained. Therefore, we hypothesized that variability in thermal preference would be higher in locales where the bet-hedging strategy is predicted to be advantageous. We established isofemale lines from gravid females sampled from seven locations across the USA, including the three used for weekly sampling. We measured variability in thermal preference of each isofemale line as the standard deviation of individual preference scores (Fig. 3a). Using Bayesian inference on a hierarchical model which treated isofemale lines as nested within their respective location (population), we estimated posterior distributions for the variability in thermal preference of each line, as well as each location (Fig. 3b). We observed strong line-to-line differences in variability among the isofemale lines measured. This is evidence for variability being under genetic control, which is a requirement for evolution by natural selection of variability as a phenotype. The line of highest variability was collected in Pennsylvania, but variability was not correlated with the predicted bet-hedging advantage across locations (*r* = −0.04, *p* = 0.92; Fig. 3c). Since these experiments used isofemale, rather than isogenic, lines, we examined if there was a correlation between variability and genetic diversity, estimated as Watterson’s θ_s_ (Supp. Fig. 3a-c). We found no significant correlation between θ_s_ and thermal preference variability as measured at either the level of isofemale lines (*r* = 0.14, *p* = 0.55) or locations (*r* = 0.027, *p* = 0.95).

**Figure 3.**
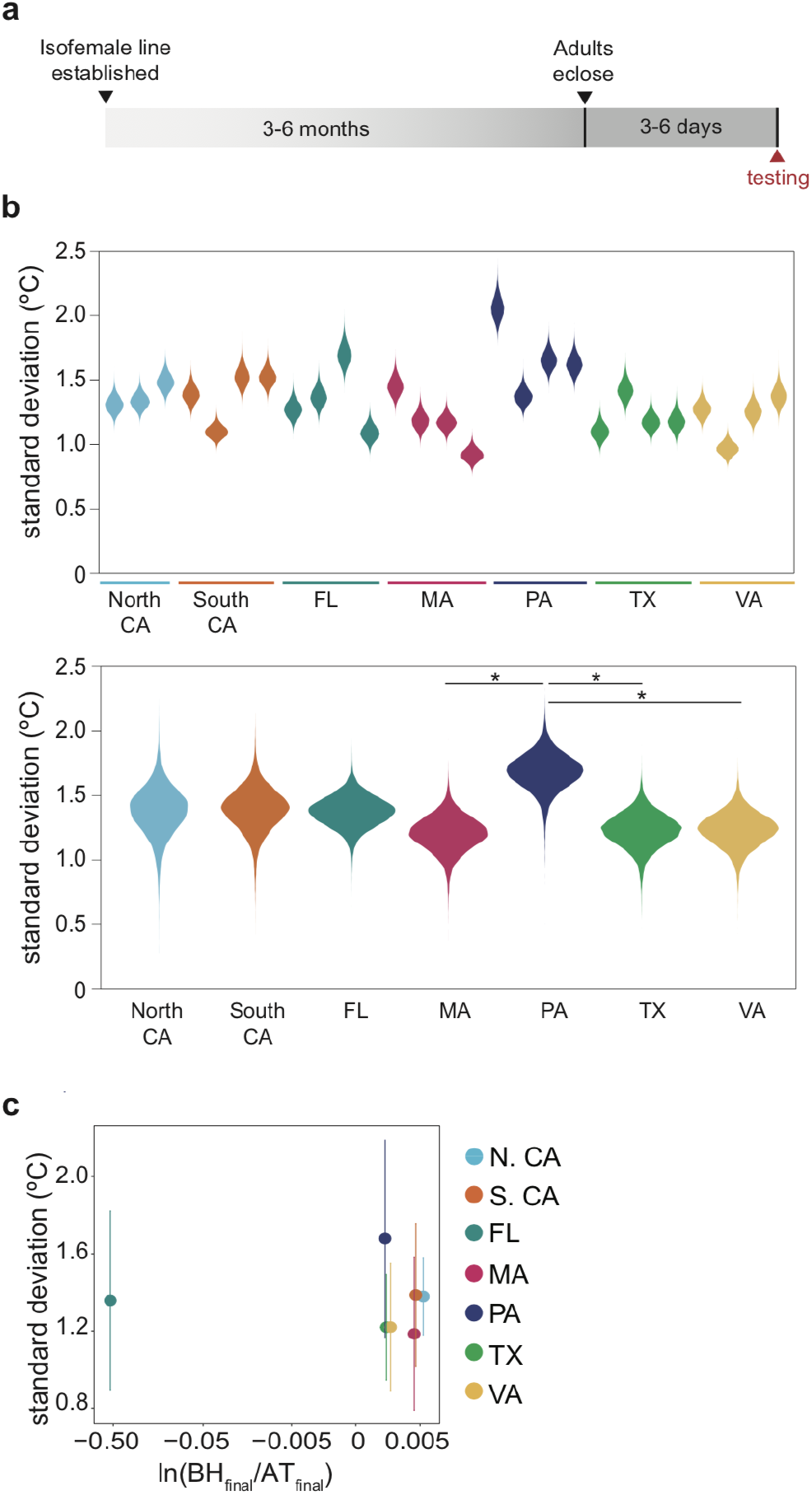
Variability of thermal preference varies among isofemale lines and is uncorrelated with predicted bet-hedging advantage. **a)** Experimental timeline for assessing variability. **b) (top)** Posterior distributions of thermal preference standard deviation for isofemale lines from each sampling location. **(bottom)** Posterior distributions of standard deviation for each sampling location. Asterisks indicate pairs of posteriors for which the 95% credible interval of the difference between locations does not include 0. See Methods for details. See Methods for details. **c)** Variability within a location vs. the bet-hedging advantage prediction for that location. Vertical error bars show the 95% credible interval of the variability posterior distribution.

### Geographic variation in heritability of thermal preference is consistent with predicted bet-hedging advantage

As a final test of the predictions of the bet-hedging model, we examined the heritability of individual thermal preferences. In bet-hedging populations, phenotypic variation does not arise due to genetic variation. Therefore, we hypothesized that heritability of thermal preference would be higher in locations predicted to favor adaptive tracking. We used isofemale lines from six locations to perform midparent-offspring regression, which measures narrow-sense heritability (*h^2^*)^41^ (Fig. 4a). We found the highest heritability in flies from FL (*h^2^* = 0.50, 95% CI = 0.24-0.75; Fig. 4b). Across all sites, *h^2^* was inversely correlated with the predicted bet-hedging advantage of the geographic origin; sites predicted to favor adaptive tracking had higher thermal preference heritability than sites predicted to favor bet-hedging (*r* = −0.90, *p* = 0.011; Fig. 4c). Removing the FL data point still produced a negative correlation, though the magnitude was smaller and the correlation was no longer significant under an α = 0.05 threshold (*r* = −0.75, *p* = 0.14). There was no significant correlation of *h^2^* with genetic diversity (Watterson’s θ_s_) (*r* = 0.54, *p* = 0.27; see Methods), but there was a significant positive correlation using PoPoolation θ_s_ estimate (*r* = 0.90, *p* = 0.015; see Supp. Fig. 3d,e). The significance of the correlation between *h^2^* and θ_s_ was driven by the data point from FL (*r* = 0.33, *p* = 0.59 with that point removed).

**Figure 4.**
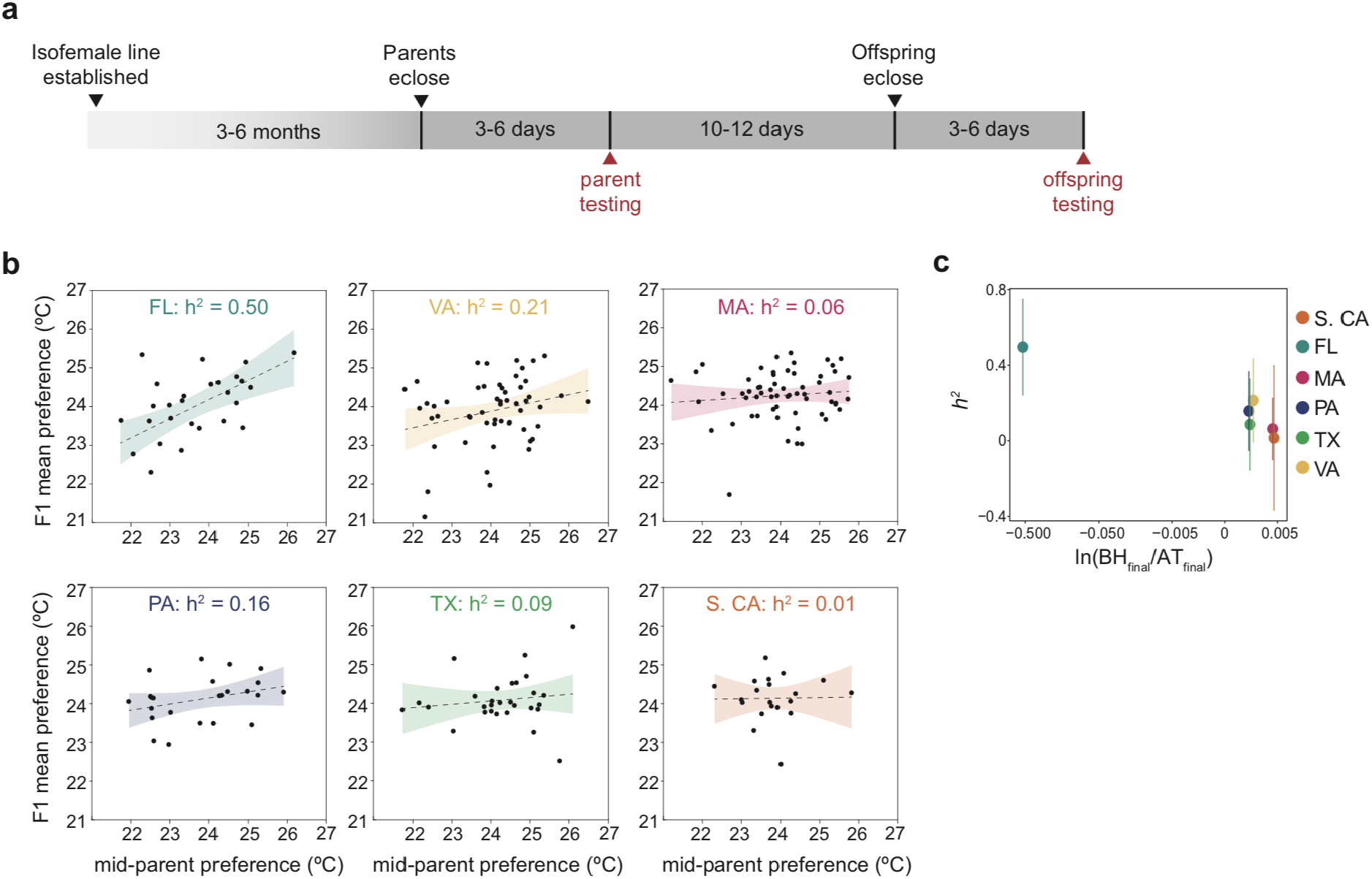
Heritability (*h^2^*) of thermal preference varies with geographic origin and is negatively correlated with the origin’s predicted bet-hedging advantage. **a)** Experimental timeline for assessing heritability. **b)** Scatter plots of F1 mean thermal preference vs midparent preference. *h^2^*, as measured by the slope of the midparent-offspring regression (dashed line; shaded area is 95% CI), varies with geographic origin of the isofemale lines. FL had the only slope that was significantly different from zero (*p* = 4.1×10^−4^), though VA was close to significant (*p* = 0.057). **c)** Scatter plot of *h^2^* vs predicted bet-hedging advantage by location. Vertical error bars show the 95% CI.

## Discussion

Bet-hedging, in which a single genotype produces a distribution of phenotypes to avoid sharp declines in fitness in the face of fluctuating selection, is a potential explanation of observed intragenotypic behavioral variability^16,32^. While this framework has strong theoretical foundations, empirical evidence, particularly in animals and with respect to behavioral phenotypes, is lacking. The goal of our study was to assess whether thermal preference variability in *Drosophila melanogaster* reflects a bet-hedging or adaptive-tracking strategy, and whether this relationship varies regionally. Overall, we found multiple pieces of evidence in favor of bet-hedging: stable mean thermal preference dynamics, substantial individuality in thermal preference, and negligible heritability of thermal preference (except in regions predicted to favor adaptive tracking). Therefore, we conclude that bet-hedging is a likely explanation of behavioral variability.

Flies have persistent individual thermal preferences that do not appear to arise from genetic or macro-environmental differences. This is consistent with the hypothesis that bet-hedging underlies preference variability, but also consistent with other mechanisms such as adaptive tracking and maladaptive developmental stochasticity. Modeling seasonal population dynamics of purely bet-hedging and adaptive-tracking fly populations using local climate data across the USA, we predicted that bet-hedging would be more successful than adaptive tracking at a large majority of sites, but with significant regional variation. This model was rooted in empirical relationships between life-history traits and thermal experience, with thermal experience determined by a combination of thermal preference and daily weather.

Given our predictions of regional differences in strategy, we collected wild flies across space (regional sites) and time (weekly at three focal sites) to test for signatures of a bet-hedging strategy in 1) dynamics of mean thermal preference, 2) variability in thermal preference, and 3) heritability of thermal preference. We found that temporal patterns in mean preference over a 20-week period for MA, VA, and FL sampling were more consistent with a bet-hedging strategy than an adaptive-tracking strategy. This aligned with our life-history model bet-hedging predictions for MA and VA, but not FL. Variability in thermal preference varied by site (consistent with being an evolvable trait), but was not correlated with predicted bet-hedging advantage. Finally, we found that heritability of thermal preference varied by site, in a pattern consistent with our bet-hedging modeling. Low thermal preference heritability was observed in the populations collected from sites predicted to be favored for bet-hedging, while the highest heritability was observed in FL flies predicted to favor adaptive tracking (Fig. 4b,c).

We found little difference among sites in thermal preference variability. However, we observed a high level of heterogeneity in thermal preference variability across isofemale lines. Differences in variability across lines suggests a genetic basis to variability, previously noted in locomotor bias^28^, phototaxis^30^, and odor preference^31^. A genetic basis for variability may be indicative of a bet-hedging trait, as optimal levels of variability could be selected for^19,21–24^. Interestingly, we also observed that rearing temperature caused plasticity in the variability of an isofemale line (Supp. Fig. 2b). We previously found plasticity in variability of locomotion and phototaxis behaviors when comparing flies that were raised in standard and enriched food vials^42^, and switching files from higher to lower quality media can also increase variability^31^. These examples of plasticity may reflect an adaptive response where stressful environmental changes cause a diversification of behaviors, i.e., a dynamic increase in bet-hedging. At a minimum, they show that both genetics and environment play roles in determining the degree of variability of behavioral traits. Since we identified differences in thermal preference heritability among our populations, we propose that the behavioral differences between individuals may be primarily due to stochastic microenvironmental forces in populations with lower heritability and allelic variation in populations with higher heritability.

In a 2011 paper, Simons established six categories of evidence for inferring the existence of a bet-hedging trait^12^. Our work integrates modeling and empirical evidence that spans all six of Simon’s levels. Evidence from the first three categories 1) identifies a potential bet-hedging trait, 2) identifies relevant environmental fluctuations, and 3) establishes genotype-level differences in phenotypic variability. We identified thermal preference as a candidate bet-hedging trait, identified seasonal temperature variation as a relevant environmental fluctuation, and showed empirically that thermal preference variability is genetically determined. Evidence from the last three categories 4) establishes that fitness varies between environments, 5) specifically when environments fluctuate, and 6) establishes that there is a quantitative match between fitness effects and fluctuating selection. Our life-history model links temperature experience/preference to fitness-affecting life-history traits via experimental measurements. Results of this model show that bet-hedging is advantageous specifically when temperatures fluctuate on timescales similar to the generation time. Finally, the model provides quantitative predictions of the degree of bet-hedging advantage across regions with different temperature fluctuations. Consistent with these predictions, we found that thermal preference heritability decreases with predicted bet-hedging advantage across sites. Through a combination of modeling and experimental measurements, we have identified a range of evidence, predominantly consistent with the bet-hedging hypothesis, that spans all of Simon’s categories.

Overall, this study provides evidence for bet-hedging in thermal preference, showing 1) high levels of non-genetic individual differences within lines and a genetic basis for the degree of variability across lines, 2) seasonal mean preference patterns consistent with bet-hedging, and 3) generally low trait heritability. Strikingly, flies in which heritability was measurably positive came from regions where adaptive tracking was predicted to be advantageous over bet-hedging. Our findings put behavioral individuality into an ecological and evolutionary context: variation that first appears like idiosyncratic ‘noise’ may reflect an adaptive strategy for dealing with risky environments.

## Methods

### Data and analysis code

All raw data and analysis code associated with this project is archived at https://zenodo.org/record/4026736 and http://lab.debivort.org/variability-reflects-bet-hedging.

### Fly husbandry

Unless otherwise stated, all stocks and isofemale lines were maintained at 22-23°C and 45% relative humidity (RH) in temperature controlled incubators in 12L:12D conditions on a yeast, cornmeal, and dextrose media (23g yeast/L, 30g cornmeal/L, 110g dextrose/L, 6.4g agar/L, and 0.12% Tegosept).

### Thermal preference assay

A two-choice assay was created to measure thermal preference, where the time-averaged temperature a fly experienced as it moved through a linear arena was the estimate of its thermal preference. The behavioral instrument consisted of 20 arenas each of which had a warm and a cool half. There was independent control of the hot and cold temperature set points, but the two set points were identical for all arenas. Temperature in the arena halves was set via Peltier elements (Custom Thermoelectric 12711-5L31-09CQ, wired in series) and resistive temperature detectors (McMaster-Carr #6568T46) under PID control by either a commercial (AccuThermo FTC100D) or custom Arduino-based controller (see code, parts list in data repository). The floor of the arena consisted of two Peltier elements separated by a ~0.5mm air gap. This design gave the ability to precisely control temperature, as well as to switch the positions of the cold and hot sides. The Peltier elements were mounted with thermal paste to aluminum blocks through which distilled water was pumped from a circulating chiller (ThermoFisher TF2500 Recirculating Chiller). Design files for the behavioral arena and Arduino controller are available in the data and analysis code repositories. Unless otherwise stated, the set point for the cold side was 20°C and the set point for the hot side was 28°C. These set points were chosen to be within the fly’s innocuous temperature range, so as to avoid activating any noxious stimuli receptors^43^, as well as being amenable to an hours-long experiment where an excessively high temperature could lead to desiccation or an excessively low temperature could lead to cessation of movement.

To estimate individual thermal preference, single flies were placed into each tunnel and allowed to freely move for 4 hours. Their positions were monitored and recorded under reflected far-infrared lighting (940nm) in an enclosed box using the beta version of the Massively Automated Real-time GUI for Object-tracking software^44^ in MATLAB 2018a (Mathworks, Inc). Two sets of behavioral arenas were simultaneously tracked by one camera, resulting in 40 flies imaged by a single camera. Three boxes were set up to work in parallel to facilitate higher throughput.

An initial thermal preference metric ranging from 0 to 1 was calculated for each individual at the end of the trial by measuring the proportion of time spent on the hot side over the total trial time (*pref* = *T*_hot_/*T*_total_). In order to correct for any bias induced by long periods of inactivity, pauses longer than 5 minutes were filtered out from the tracks and the thermal preference was recalculated. Flies which had less than 1 hour total activity throughout the trial post-filtering were removed from further analysis. Small deviations from the hot and cold set points were observed among the different Peltier elements, so a tunnel-specific temperature correction was applied to the each fly’s 0-1 metric:

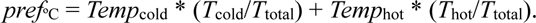

The tunnel correction gives a thermal preference metric in °C, based on the measured tunnel temperatures, which translates to the thermal experience of a fly given the time spent at the cold and hot temperatures.

### Bet-hedging and adaptive tracking model

Predictions for seasonal patterns in mean thermal preference and calculations of bet-hedging advantage were made using a modified version of a previous evolutionary model^32^. In brief, the current and original models use a system of difference equations coupled to empirically determined relationships between temperature and development time and lifespan to model fly populations over a breeding season. Simulations were performed in MATLAB 2018a (Mathworks, Inc). Thermal preference of simulated flies was determined by one of two pure strategies: bet-hedging (no heritability in thermal preference; thermal preference of the new individuals determined by sampling from a beta distribution, a close fit to the observed distribution) or adaptive tracking (heritability of 1, thermal preference of offspring is determined entirely by parents). The model incorporated first order terms for births of new flies (β) and deaths by causes other than temperature-dependent old age (δ). These were calibrated using two assumptions: 1) the initial population size matched the final population size at the end of the breeding season (the population is in steady state with the environment) and 2) the mean preference at the start of the season matched the mean preference at the end of the season (flies have adapted to local conditions). These constraints were evaluated under the adaptive tracking strategy, and the specific values of the β and δ parameters that satisfy these constraints were determined by a hill-climbing algorithm. Using climate normals (average daily temperature), a breeding season was set to start when temperatures exceed 6.5°C and end when they drop below 10°C. Normal values were smoothed with a two-month moving average in identifying the first and last days of the season.

Two aspects of the original model were updated here: 1) determination of thermal experience given a thermal preference and the available environmental temperature range and 2) empirical relationships between temperature and development time/life-span. Under the previous approach, thermal experience *τ* of fly *i* on day *j* was calculated as:

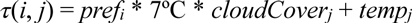

Where *pref_i_* is thermal preference of fly *i* (0-1 scale), 7°C is a typical empirical difference between sun and shade temperatures^32^, *cloudCover_j_* is the fraction of cloud cover on day *j*, and *temp_j_* is the average in-shade temperature on day *j*. This coding of thermal experience produces a 7°C difference in thermal experience between flies at the thermal preference extremes, without consideration of flies avoiding noxiously hot and cold temperatures^31^ (Supp. Fig. 4d). Under the updated approach, thermal experience was determined by a piecewise function to allow even flies with extreme thermal preferences to avoid experiencing noxious temperatures:

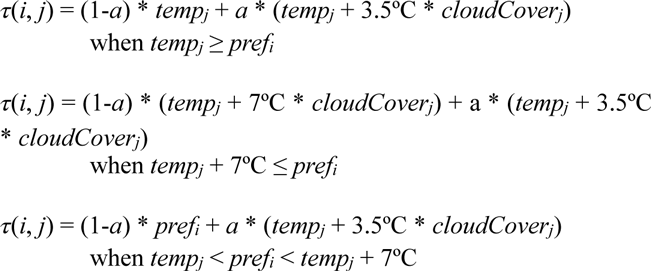

where *temp_j_* and *cloudCover_j_* are as above, *pref_i_* is thermal preference of fly *i* in °C, and *a* is a value between 0 and 1 reflecting the degree to which a fly’s thermal experience is locked in by ambient conditions. *pref_i_* ranges between 18°C and 30°C, the limits of the thermal preference assay. The new formula specifies that when the preference of fly *i* is between the in-shade and in-sun temperatures of day *j*, the fly’s thermal experience is a combination of the proportion of time spent (1-*a*) at the thermal preference temperature, *pref_i_* (reflecting thermoregulatory behavior), and proportion of time (*a*) spent at the average daily temperature, *temp_j_* + 3.5°C x *cloudCover_j_* (reflecting non-thermoregulated behaviors, such as predator avoidance or foraging). When the in-sun temperature for day *j*, *temp_j_* + 7°C x *cloudCover_j_*, is less than or equal to the fly’s thermal preference, the fly will spend the thermoregulatory portion of their behavior at the in-sun temperature (maximum temperature it can achieve). When the in-shade temperature, *temp_j_*, is greater than or equal to the fly’s thermal preference, the fly will spend the thermoregulatory portion of their behavior at the in-shade temperature (minimum temperature it can achieve). For a sun vs. shade temperature difference of 7°C, an *a* of 0.4 was chosen, which is equivalent to saying that when balanced against other behavioral demands, flies are able to achieve 60% of their desired thermoregulation. This value was determined by matching the standard deviation of thermal experience over the entire breeding season of simulated flies to ~1.5°C, the average measured standard deviation in the laboratory thermal preference assay (Supp. Fig. 4, Fig. 3b). The relative bet-hedging advantages calculated with the model are robust to different *a* values (Supp. Table 1). To create the map of bet-hedging advantage across the USA, simulated fly thermal preferences were drawn from a beta distribution with μ = 0.44 and σ^2^ = 0.015 for all stations, with the mean and variance representing a typical thermal preference behavioral statistics of a wild fly population.

The relationships of development time and lifespan to thermal experience were updated based on data collected from three isofemale lines from Coral Gables, Florida, USA (FL), Cambridge, Massachusetts, USA (MA), and Charlottesville, Virginia, USA (VA) (Supp. Fig. 5). To parameterize the relationship between development time and temperature, a quadratic fit on combined data from the three isofemale lines was used (trends from individual lines were similar). For lifespan vs. temperature, a natural logarithm fit to the combined data was used. For fly *i* and on day *j*, development time and lifespan are determined by the following equations:

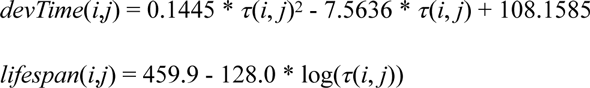

where *τ*(*i, j*) is the thermal experience of fly *i* on day *j*.

### Seasonal measurement of mean thermal preference

Wild flies were collected weekly from June 24, 2018 until November 1, 2018 in Cambridge, Massachusetts, USA, Charlottesville, Virginia, USA, and Coral Gables, Florida, USA In VA, *Drosophila* were collected via aspiration from rotting fruit at Carter Mountain Orchard (37.99° N, 78.47° W). In MA and FL, baited traps were set out to capture *Drosophila* around the residential areas in the vicinity of Harvard University (42.38° N, 71.12° W) and University of Miami (25.72° N, 80.28° W).

The MA and FL traps were created by cutting an approximately 2 inch-square flap into an empty one-gallon plastic ethanol jug (Koptec, Decon Labs) and baiting them with a fruit and wine mixture. Flies were collected by placing an empty fly food bottle over the neck of the jug and allowing flies in the trap to move upwards into the bottle. Bait for each trap consisted of two bananas sliced ~7mm thick and an orange, cut in half, soaked overnight in 50mL of 8.5% alc/vol red wine. Bait was added to the trap, sprinkled with dried baker’s yeast grains, and the trap was hung on a fence or railing 2-3 days prior to the start of the week’s collections. At the end of the week’s collections, the trap was removed and thoroughly washed to get rid of any bait/larvae/pupae before fresh bait was put back in.

Collected flies were taken back to the local lab and sorted by sex and species (see Supp. Fig. 6 for detailed experimental timeline for each location). For the thermal preference assay, *D. melanogaster* males and *D. melanogaster/D. simulans* females were chosen (visual species identification of female *melanogaster* and *simulans* was too difficult to perform at scale). Female flies were housed individually after the thermal preference assay. Species identification was performed on their male offspring. Females that did not produce an F1 generation were identified through sequencing of *CoII* gene (forward primer: 5’ - ATGGCAGATTAGTxGCAATGG; reverse primer: 5’-GTT-TAAGAGACCAGTACTTG).

Flies caught in MA were assayed immediately after being collected. VA and FL females were housed in vials for 1-7 days post-collection to collect eggs prior to behavior assaying (due to higher lethality in the behavior assay at these sites). Caught flies were stored in vials at 22-23°C in ambient laboratory conditions. A mean thermal preference was determined from all the flies sampled on a particular week.

The consistency of observed mean preferences over the 20 week sampling period with predicted dynamics of purely bet-hedging or adaptive-tracking populations was assessed using the log-likelihood ratio. Predicted dynamics were calculated using location-specific 2018 daily average temperatures^38^ plugged into the lifehistory model. The mean and variance parameters used to sample fly thermal preferences in the model were determined in a location-specific manner using the assayed flies from each location: MA: μ = 0.45, σ^2^ = 0.016, VA: μ = 0.37, σ^2^ = 0.019, FL: μ = 0.49, σ^2^ = 0.010. Birth (β) and death (δ) rates were calibrated for each population independently using NOAA daily average 2018 temperatures for MA and VA and NOAA climate normals for FL (due to failure of calibration on 2018 daily temperatures). The log-likelihood of the bet-hedging and adaptive-tracking models given the observed mean preferences was calculated as:

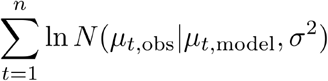

where *n* is the number of weeks of data, *N* is the normal distribution probability density function, ***μ***_*t*,obs_ is the observed mean preference on a particular week for a particular site, ***μ***_*t*,model_ is the predicted mean preference under either model on a particular week for a particular site, and *σ*^2^ is the empirical variance of thermal preference measured for flies from that site. Preference data from individual flies was bootstrapped on a week-by-week basis 1000 times to determine the error in the estimate of the log likelihood ratio between the bet-hedging and adaptive-tracking models. The findings are robust to different *a* values, as well as different parameterizations of the relationship between temperature and life-history (Supp. Fig. 7).

### Thermal preference variability of geographically diverse lines

Wild gravid *D. melanogaster* females caught in Houston, Texas, USA (TX; 29.76° N, 95.36° W) and Oakland, California, USA (Northern CA; 37.80° N, 122.27° W) in September 2018 and Pasadena, California, USA (Southern CA; 34.15° N, 118.14° W) and Media, Pennsylvania, USA (PA; 39.89° N, 75.41° W) in October 2018 were used to establish isofemale lines. In addition, using animals from the seasonal collection experiment, many isofemale lines were established from flies collected in FL, MA, and VA from July to September 2018. Four isofemale lines from each location (with the exception of Berkeley, where there were three) were chosen randomly for evaluation of thermal preference variability.

200-250 mated female flies (aged 3-6 days) from FL, MA, TX, and VA isofemale lines were assayed for thermal preference in two batches (two lines from each location per batch) in November and December 2018. Northern CA, Southern CA, and PA flies were tested in two batches in late January 2019 and April 2019. Two MA isofemale lines previously tested earlier were retested alongside these lines to control for batch effects on variability. Batch effects were calculated as the average difference in variability for the internal control lines between the November/December 2018 trials and the January 2019 or April 2019 trials. Batch effects were added to the variance estimates of the isofemale lines to remove the effect of the time of year on variability.

Bayesian inference was used on a hierarchical model to estimate the mean and variance of thermal preference (in °C) in each isofemale line and for each location. In the hierarchical model, isofemale lines were nested within sampling location, such that the prior on the line mean and variance was dependent on the hyper-prior for the location:

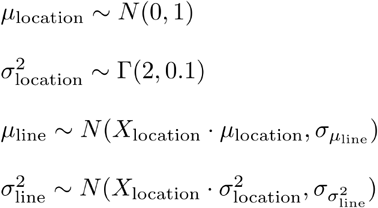

The likelihood was specified as follows:

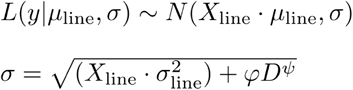

where *y* is the vector of observed thermal preferences (°C), *X* is dummy-coded predictor matrix for either line or location categories, and *D* is a vector of distance traveled during the experiment. *σ* depends on both the line variance (*σ*^2^_line_) and a sampling error component (*φD^ψ^*) that depends on distance traveled during the experiment (there is more noise in the estimate of thermal preference of flies that do not move much in the assay)^31^. *φ* and *ψ* constants were calculated by fitting a power function to the relationship between variance and distance traveled for flies experiencing no temperature stimulus. This allows us to differentiate variance that is inherent to the line from variance that comes from sampling noise due to variable activity in the assay.

Bayesian inference was done using R’s Stan interface v.2.18.2^45^. Posterior distributions for mean and variance for both lines and locations were generated by sampling using four chains: 25000 iterations per chain, with target average proposal acceptance probability of 0.9, and maximum tree depth of 10. Each chain’s effective sample size was determined to be ~12000. Every other sample from the chain was saved to the posterior distribution to reduce autocorrelation between the samples. As a measure of variability, we used the standard deviation, which was calculated by taking the square root of the variance at each step in the chain. Model fits were qualitatively evaluated using graphical posterior predictive checks, where mock data that were generated using values from the posterior distributions were compared to our observed data.

To establish whether variability estimates between two locations were different from each other, we generated the posterior distribution of differences by subtracting variability estimates for one location from the other at each step in the chain. If the 95% credible interval for the distribution of differences did not include 0, the two locations were considered to be different from each other in terms of variability^42,46^.

### Heritability of thermal preference

Narrow-sense heritability was calculated using parent-offspring regression. Males and females from 5 isofemale lines from the Southern CA, PA, and TX sampling locations were chosen for the parental generation. Males and females from 10 isofemale lines from FL, MA, and VA sampling locations were chosen for the parental generation for these sites. Parents were collected, separated by sex to maintain virginity of females, and aged 3-6 days before testing for thermal preference. After testing, flies were housed individually. 10 crosses were made from the Southern CA, PA, and TX parents, and 20 crosses were made from the FL, MA, and VA parents (Supp. Fig. 8). Each cross between two lines was replicated 2-3 times with independent sets of parents as backups in case some crosses did not produce progeny. Parent flies had to pass the activity thresholds for the thermal preference assay to be included in the crossing scheme.

Male F1s from each cross were collected and aged 3-7 days prior to testing. For each cross, a minimum of four male F1s were tested, with 10 male F1s tested for 96% (221/230) of crosses. All crosses with fewer than three male F1s passing the thermal preference filtering threshold were excluded from further analysis. Narrow-sense heritability was estimated from the slope of the regression of F1 mean thermal preference on the mid-parent thermal preference.

### Plasticity in thermal preference

Males and females (0-2 days old) were collected from a MA isofemale established in mid-August and allowed to lay eggs for 48 hours at 26°C (45% RH) and 22°C (45% RH) and 96 hours at 18°C (50-55% RH). Female F1s were collected daily and aged 3-6 days with males (to ensure mated status) at the treatment temperature prior to thermal preference testing. Mean and variability in thermal preference was estimated using Bayesian inference in Stan, as above. The priors and likelihood function were as follows:

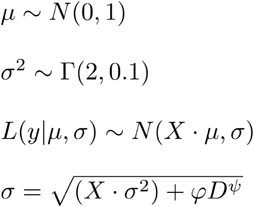

where *y* is the vector of observed thermal preferences, *X* is dummy-coded predictor matrix for the temperature treatments, and *D* is a vector of distance traveled during the experiment. As with the variability experiments, *σ* is partitioned into the line variance (*σ*^2^) and sampling noise (*φD*^ψ^).

### Estimating genetic diversity in the sampled populations

Genomic DNA from 4 female flies from each isofemale line tested in the heritability and variability assays was extracted using bead-beating and the ZYMO *Quick*-DNA kit (cat. no. D3012). DNA was made into libraries using a liquid handling robot (Analytic Jena CyBio-Felix Model 30-5015-100-24). Library preparation was done using a tagmentation protocol with Tn5 transposase^47,48^. Genomic DNA from 273 individual flies was made into per-individual libraries. 150bp paired-end reads were sequenced on an Illumina NovaSeq platform with mean 0.02x - 8x coverage per individual. Alignment of reads was done using the BWA-MEM algorithm (v0.7.15; default parameters)^49^ to the *Drosophila melanogaster* reference genome 6.28 release. PCR and optical duplicates were flagged using Picard’s Mark-Duplicates (v2.20.6). HaplotypeCaller in GATK version 4.1.3.0 was used to call variants^50^. Given the low sequencing coverage, minimum pruning support and minimum dangling branch length were set to 1; all other parameters were kept at default values. Mean coverage depth and fraction of missing genotypes per individual was quantified using VCFtools^51^. Individuals with mean coverage depth less than 2x were excluded from further analysis, leaving a total of 246 individuals.

Variants were filtered for biallelic SNPs with a minor allele frequency > 5%. 15,080 SNPs (distributed across the genome) with called genotypes for all individuals were used to calculate Watterson’s θ_s_. A genotype matrix of variants by individuals was created. Cells of this matrix indicate whether a particular individual is homozygous for the reference or alternate allele or heterozygous. Since the number of individuals and number of lines in each population will influence the θ_s_ estimate, the subsampling approach described below was used.

Individuals were divided into subsets: those used in the heritability analysis and those used in the variability analysis. For each subset, the geographic population that had the fewest lines and individuals was chosen as the subsampling benchmark, and all other populations were subsampled to that benchmark size. To estimate θ_s_ in a population and its uncertainty, a bootstrapping approach was employed in addition to the subsampling. For each bootstrap iteration, individuals from the target population that had the same number of lines and the same number of individuals per line as the benchmark population were sampled with replacement. For each variant, a homozygous individual would contribute either a reference or alternate allele, and a heterozygous individual would contribute a reference allele with a 50% probability. A variant was counted as segregating if individuals contributed both reference and alternate alleles. The number of segregating sites and Watterson’s theta (θ_s_) were then calculated from the chosen individuals:

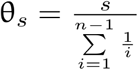

where *S* is the number of segregating sites and *n* is the number of individuals. The final result was a metric of genetic diversity that could be compared across populations of different individual and line compositions (Supp. Fig. 9).

To calculate within-line genetic diversity, the same general approach as above for calculating θ_s_ was employed. Since there was a maximum of four individuals per line, a bootstrapping approach was not used. Instead, θ_s_ was calculated only for lines that had the full complement of four individuals (Supp. Fig. 9).

As a complement to the bootstrapping approach, θ_s_ was also calculated using PoPoolation^40^ for both the heritability and variability flies. Using only the 246 individuals with mean coverage > 2x, the number of reads per individual was downsampled using SAMtools^52^ to match the individual with the lowest coverage in order to standardize coverage across individuals. As in the bootstrapping approach, populations with more isofemale lines were subsampled to match the population with the fewest isofemale lines prior to making the population pileup file. The pileup file was then filtered using the *identify-genomic-indel-regions. pl* and *filter-pileup-by-gtf. pl* functions to remove indels and the variants within 5bp of them. θ_s_ was calculated from the pileup file in 50kb non-overlapping windows using only SNPs with a minimum minor allele count of 2, minimum site coverage of 4, maximum site coverage of 400, and reads with a minimum quality score of 20.60% of the 50kb window had to have coverage between 4 and 400 for θ_s_ to be calculated. To get a single population θ_s_, the mean of θ_s_ across all 50kb windows for chromosomes 2L, 2R, 3L, 3R, X, and 4 was taken (Supp. Fig. 9). The θ_s_ from the PoPoolation analysis is reported as θ_s_/nt to distinguish it from our bootstrapping approach.

## Acknowledgements

We thank Nick Keiser, Alex Keene, Sophie Caron, John Tuthill, and Rob Unckless for kindly supplying wild-derived isofemale lines. Brian J. Arnold and Luisa Pallares provided invaluable assistance with sequencing data analysis and Elena Filippova provided expert help with genomic library preparation. We thank Edward Soucy and Brett Graham of the Center for Brain Science Neurotechnology Core for help with instrument manufacturing and design. JAZ was supported by The NSF-Simons Center for Mathematical and Statistical Analysis of Biology at Harvard, award number #1764269 and the Harvard Quantitative Biology Initiative. BdB was supported by a Sloan Research Fellowship, a Klingenstein-Simons Fellowship Award, a Smith Family Odyssey Award, a Harvard/MIT Basic Neuroscience Grant, the NSF under grant no. IOS-1557913, and the NIH under grant no. MH119092.

## Supplementary figures and table

**Supp. Figure 1.**
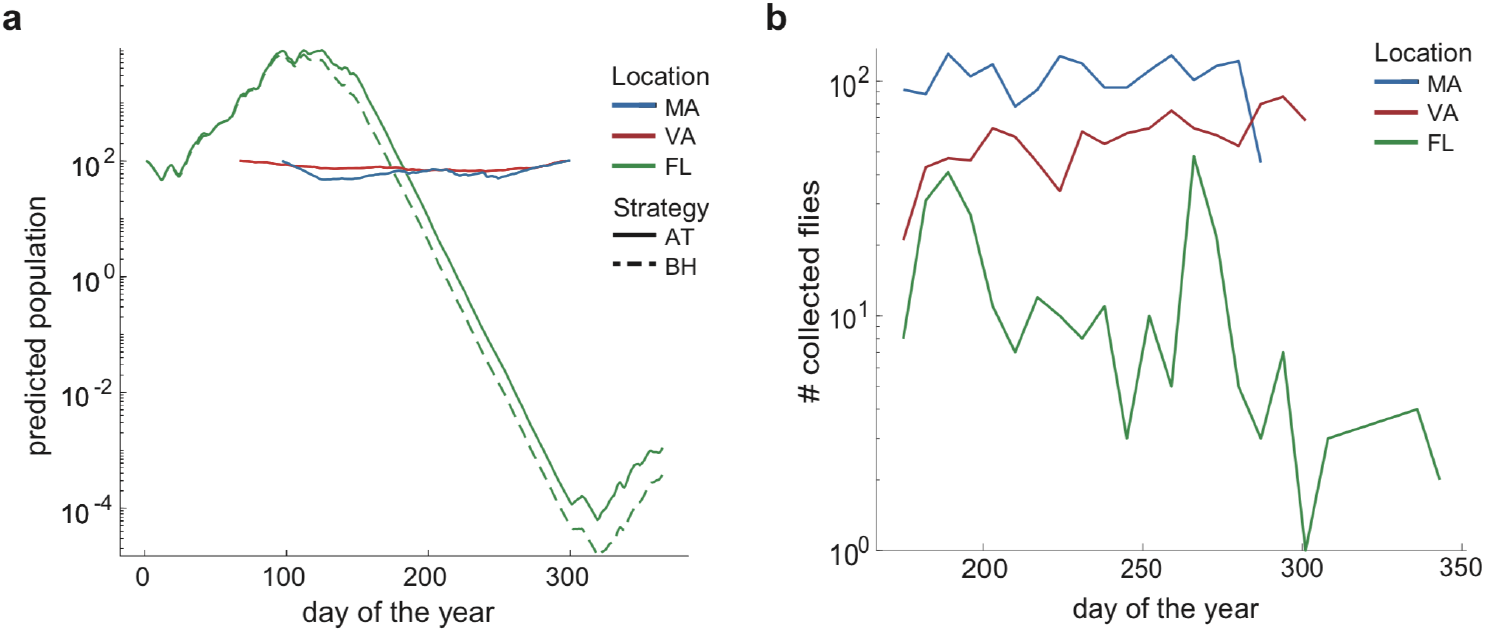
Predicted population dynamics in 2018 and numbers of collected *D. melanogaster*. **a)** Predicted population sizes under adaptive tracking (AT; solid) and bet-hedging (BH; dashed) strategies for the 2018 breeding season in Cambridge, Massachusetts, USA (MA), Charlottesville, Virginia, USA (VA), and Coral Gables, Florida, USA (FL). The predicted differences between population sizes for AT and BH strategies in MA and VA overlap on this scale. **b)** Logged number of collected *D. melanogaster* over the 2018 collection period across the three sampling locations. The timing of the weeks with fewest flies collected in FL coincide with the model’s lowest predicted population size.

**Supp. Figure 2.**
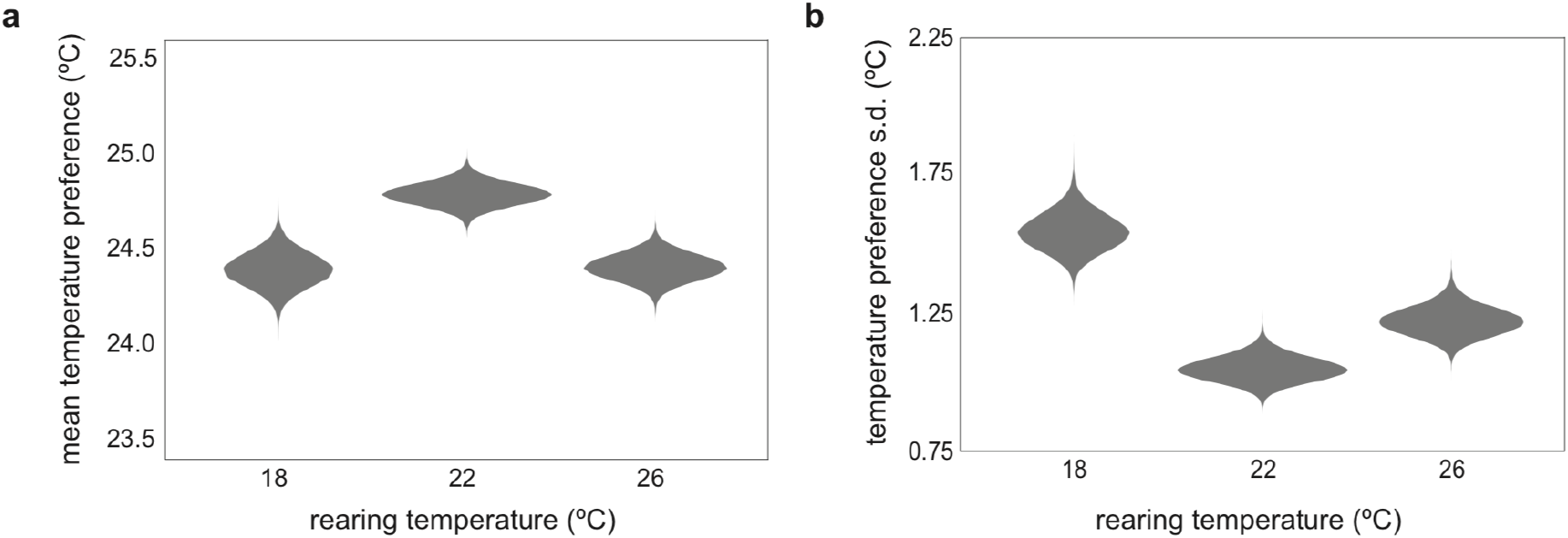
Plasticity in mean and standard deviation of thermal preference. **a)** Posterior estimates of mean temperature preference under three different rearing temperatures (18°C: *n* = 134; 22°C: *n* = 166; 26°C: *n* = 145). **b)** Posterior estimates of variability (s. d.) under three different rearing temperatures.

**Supp. Figure 3.**
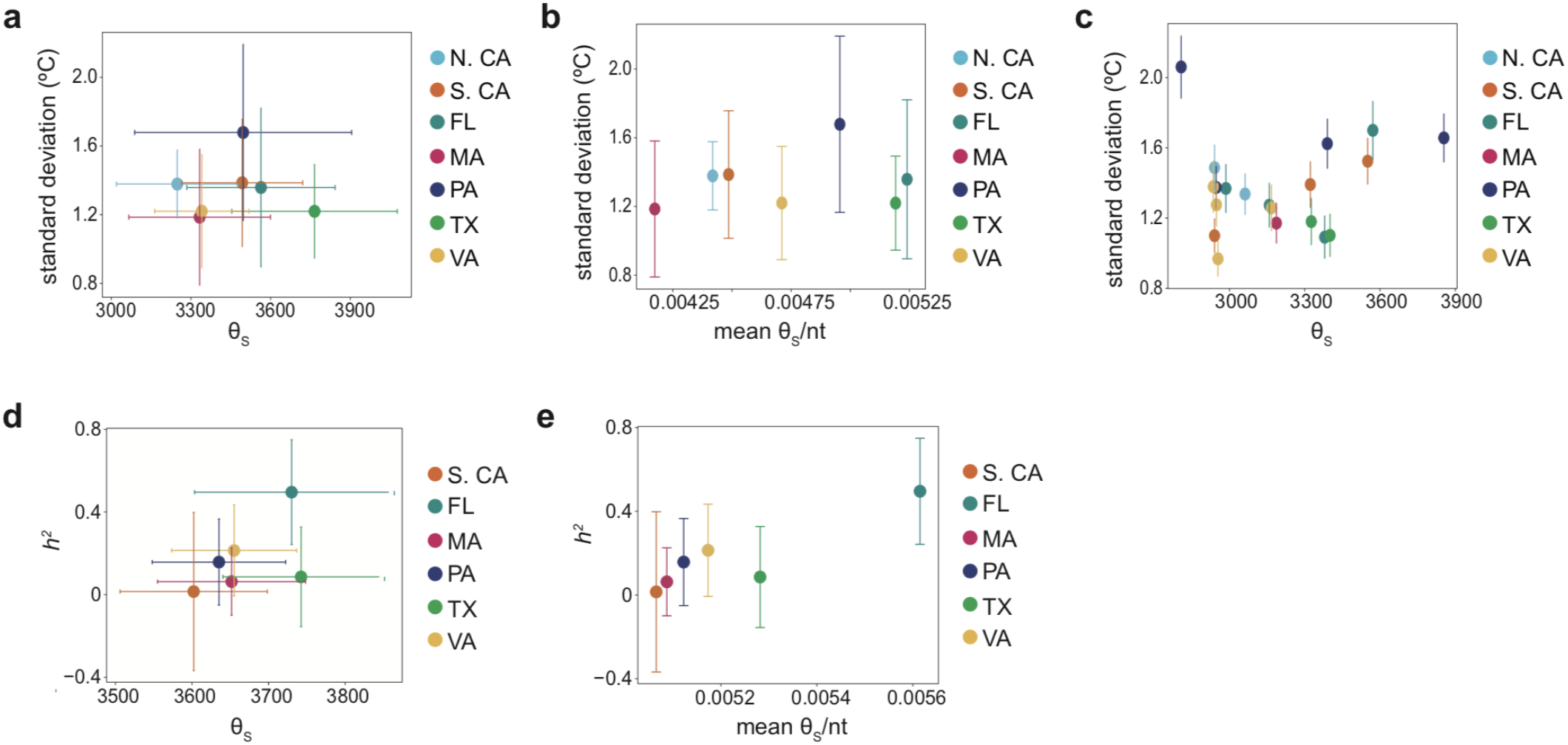
Genetic diversity as a potential predictor of thermal preference variability and heritability. **a)** θ_s_ within a location vs. the variability estimate for that location (*n* = 7; *r* = 0.027, *p* = 0.95). Vertical error bars show +/− 2 s. d. of the variability posterior distribution and horizontal error bars show +/− 2 s. d. of the bootstrapped distribution of θ_s_. **b)** Average PoPoolation^40^ estimate of θ_s_/nt within a location vs. thermal preference variability (*r* = 0.23, *p* = 0.62). **c)** θ_s_ within an isofemale line vs. thermal preference variability for that line (*n* = 20; *r* = 0.14, *p* = 0.55). Vertical error bars show +/− 2 s. d. of the variability posterior distribution. **d)** Site-specific heritability (*h^2^*) vs θ_s_ (*n* = 6; *r* = 0.54, *p* = 0.27). Vertical error bars show the 95% CI on the *h^2^* estimate (regression slope) and horizontal error bars show +/− 2 s. d. of the bootstrapped distribution of θ_s_. **e)** *h^2^* vs Average PoPoolation estimate of θ_s_/nt by location (*r* = 0.90, *p* = 0.015). Removing the FL data point diminished the positive correlation (*r* = 0.33, *p* = 0.59).

**Supp. Figure 4.**
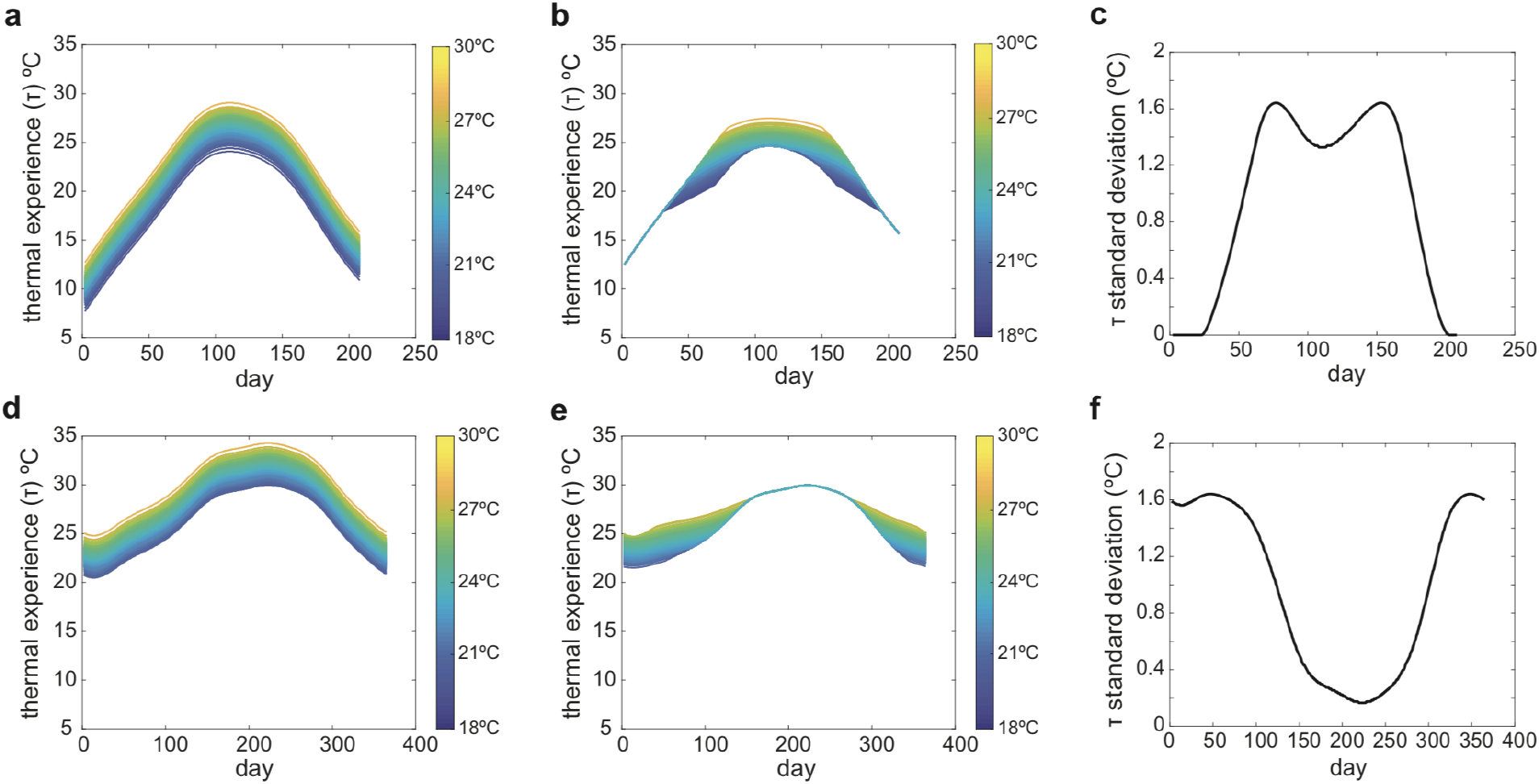
Relationships between thermal preference and thermal experience (*τ*) in the original and updated models. Colored lines in a-b) and d-e) reflect flies with particular thermal preferences bounded by 18°C and 30°C (color scale). a) *τ* under an average Boston, Massachusetts, USA breeding season in the original model. b) *τ* under an average Boston breeding season in the updated model. c) Standard deviation in *τ* over the Boston breeding season for the updated model. d) *τ* under an average Miami, Florida, USA breeding season in the original model. e) *τ* under an average Miami breeding season in the updated model. f) Standard deviation in *τ* over the Miami breeding season for the updated model.

**Supp. Table 1.** Bet-hedging advantage under different *a* values. Bet-hedging advantage was calculated as ln(BH_pop, final_/AT_pop, final_). Station-specific climate normals were used to calculate the bet-hedging advantage (behavioral parameters: μ = 0.44, σ^2^ = 0.015). A dash (-) signifies that the β and δ parameters could not be calibrated and no estimate of bet-hedging advantage was calculated. Bet-hedging advantage increases as flies are more able to implement their thermal preference (lower *a*), but the qualitative pattern of bet-hedging advantage across sites is robust to the choice of *a*.

| Location (Station ID) | $a = 0.7$ | $a = 0.5$ | $a = 0.4$ | $a = 0.3$ | $a = 0.2$ |
| --- | --- | --- | --- | --- | --- |
| Berkeley, California, USA (USC00040693) | $8.6 \times 10^{-4}$ | $2.6 \times 10^{-3}$ | $3.7 \times 10^{-3}$ | $5.3 \times 10^{-3}$ | $7.0 \times 10^{-3}$ |
| Boston, Massachusetts, USA (USW00014739) | $1.3 \times 10^{-3}$ | $2.9 \times 10^{-3}$ | $3.8 \times 10^{-3}$ | $4.7 \times 10^{-3}$ | $5.7 \times 10^{-3}$ |
| Philadelphia, Pennsylvania, USA (USW00013739) | $3.4 \times 10^{-4}$ | $7.2 \times 10^{-4}$ | $9.1 \times 10^{-4}$ | $1.1 \times 10^{-3}$ | $1.3 \times 10^{-3}$ |
| Charlottesville, Virginia, USA (USW00003759) | $8.1 \times 10^{-4}$ | $1.6 \times 10^{-3}$ | $1.9 \times 10^{-3}$ | $2.4 \times 10^{-3}$ | $3.0 \times 10^{-3}$ |
| Pasadena, California, USA (USC00046719) | $3.3 \times 10^{-4}$ | $6.5 \times 10^{-4}$ | $9.9 \times 10^{-4}$ | $1.1 \times 10^{-3}$ | $1.3 \times 10^{-3}$ |
| Houston, Texas, USA (USW00012918) | $4.7 \times 10^{-3}$ | $5.4 \times 10^{-3}$ | $7.8 \times 10^{-3}$ | - | - |
| Miami, Florida, USA (USC00085667) | -0.17 | -0.35 | -0.45 | -0.57 | -0.68 |

**Supp. Figure 5.**
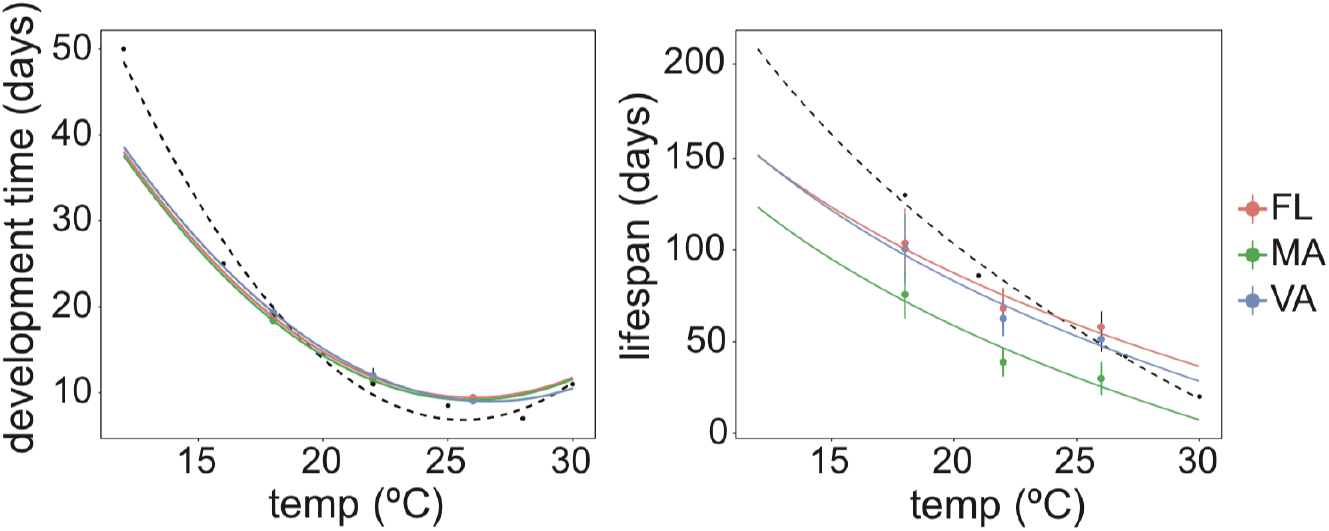
Life-history data used in the original model (black dashed lines) and updated model life-history data using isofemale lines from three locations (colored lines). Error bars show the 95% CI of the mean.

**Supp. Figure 6.**
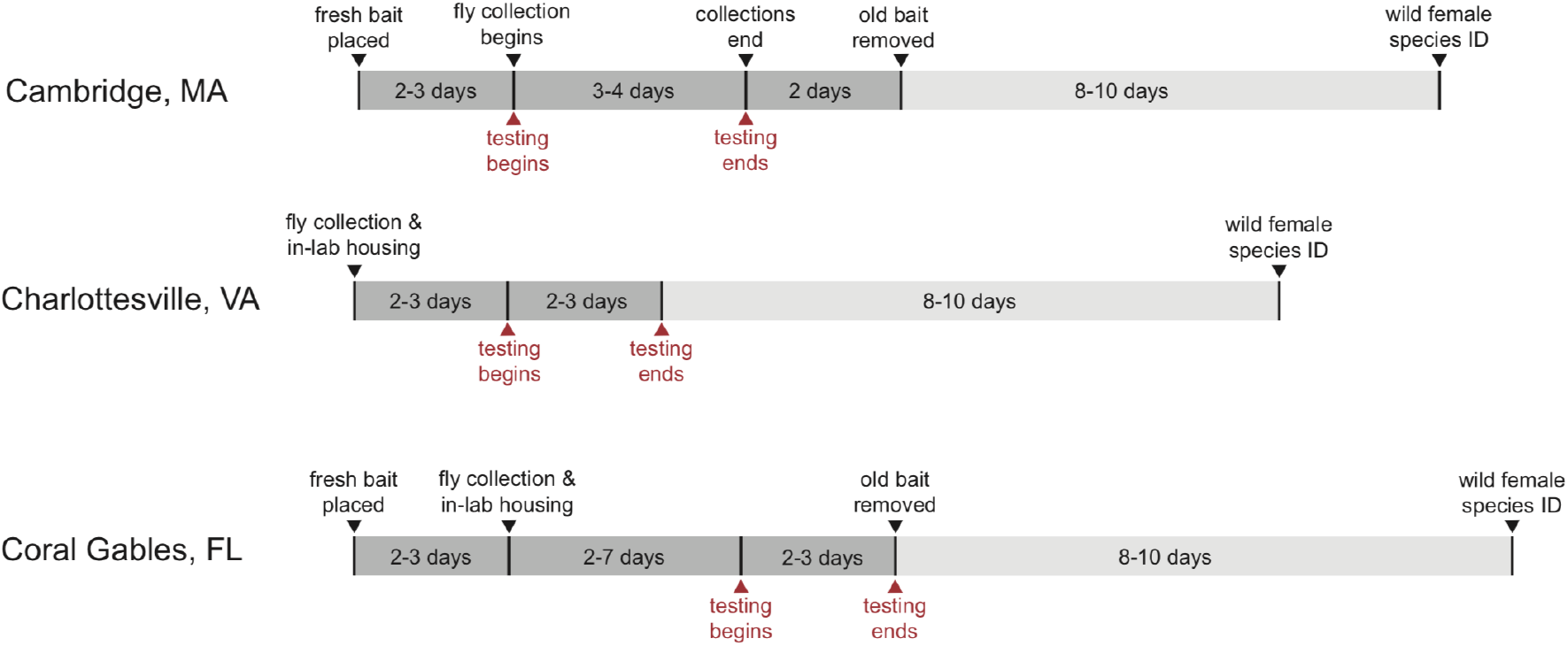
Experimental timelines for seasonal collections at all three locations. Each timeline shows the process of collection and testing for flies collected each week.

**Supp. Figure 7.**
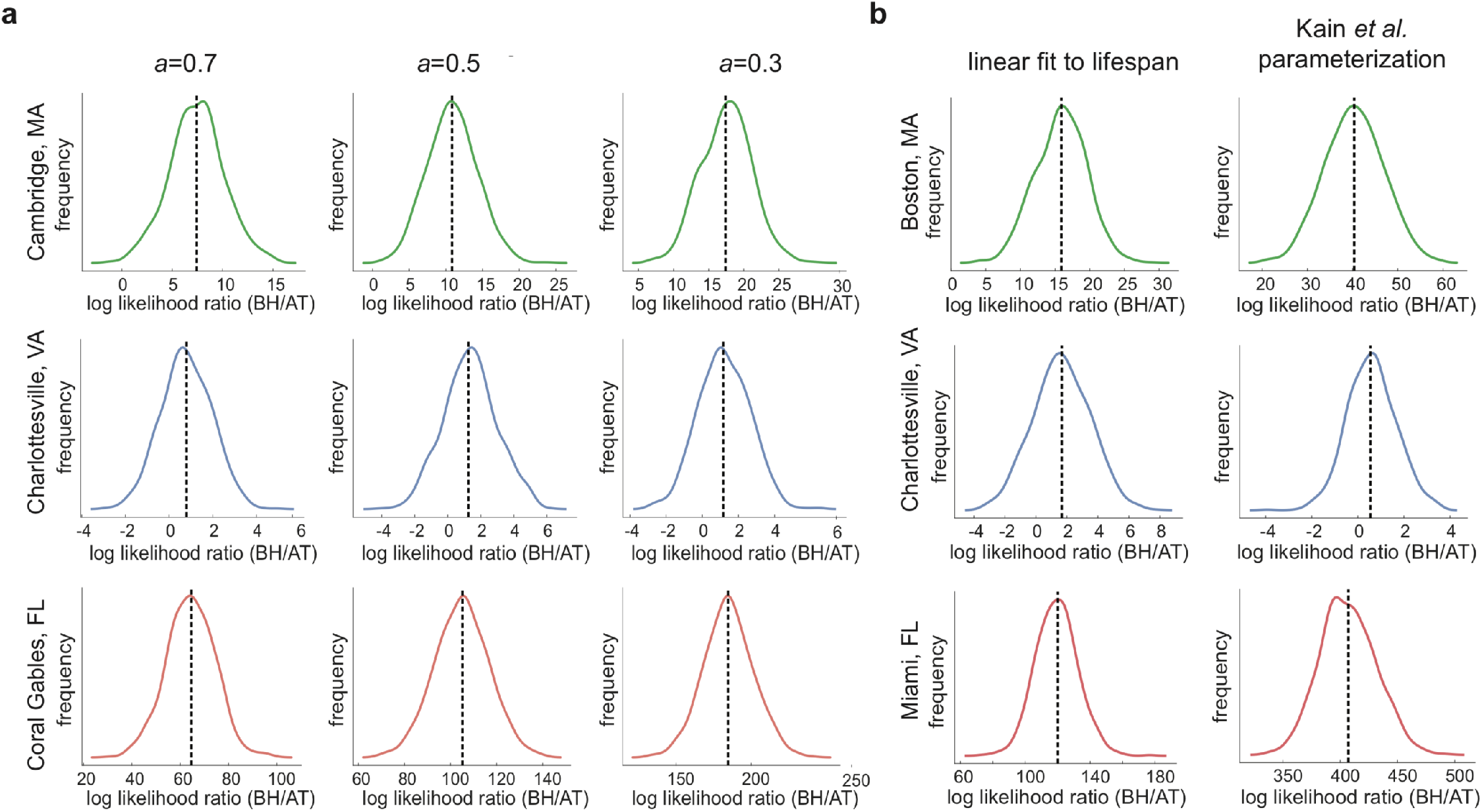
Robustness of mean preference log-likelihood ratios to the values of *a* and lifespan-temperature relationship parameterization. **a)** Kernel density estimates of the log-likelihood ratio of the (bet-hedging/adaptive-tracking) models for seasonal mean preferences, over bootstrap replicates. Columns of plots reflect different values of *a*. Dashed lines show the observed log-likelihood ratio for all the data. **b)** As in (a) for alternative parameterizations of the relationship between lifespan and temperature (*a* = 0.4).

**Supp. Figure 8.**
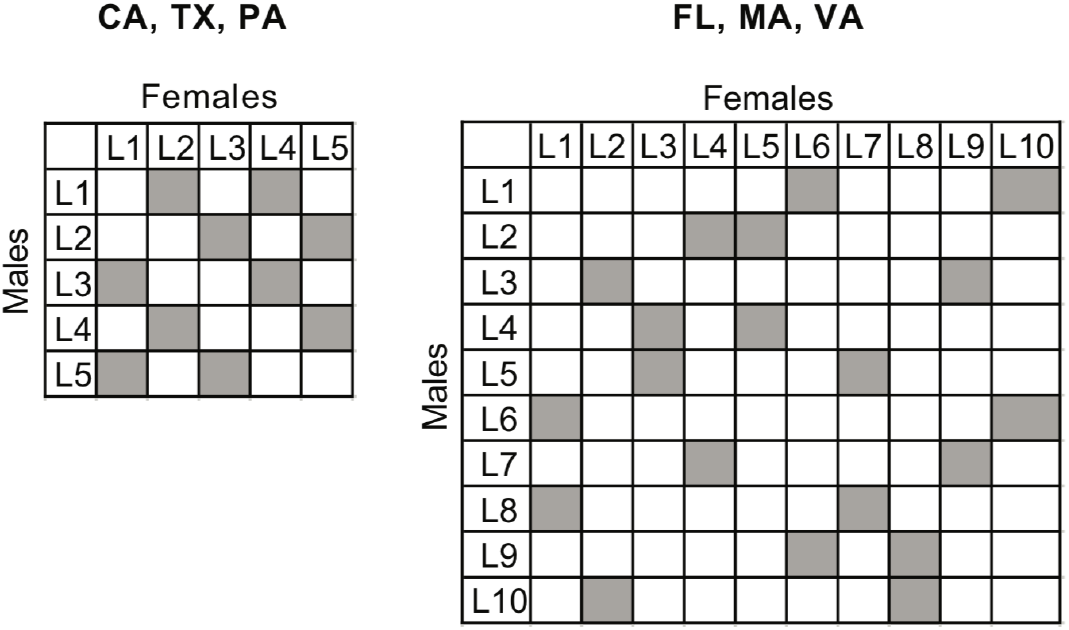
Cross scheme for heritability analysis. Gray squares indicate a cross between two lines (no crosses were made along the diagonal).

**Supp. Figure 9.**
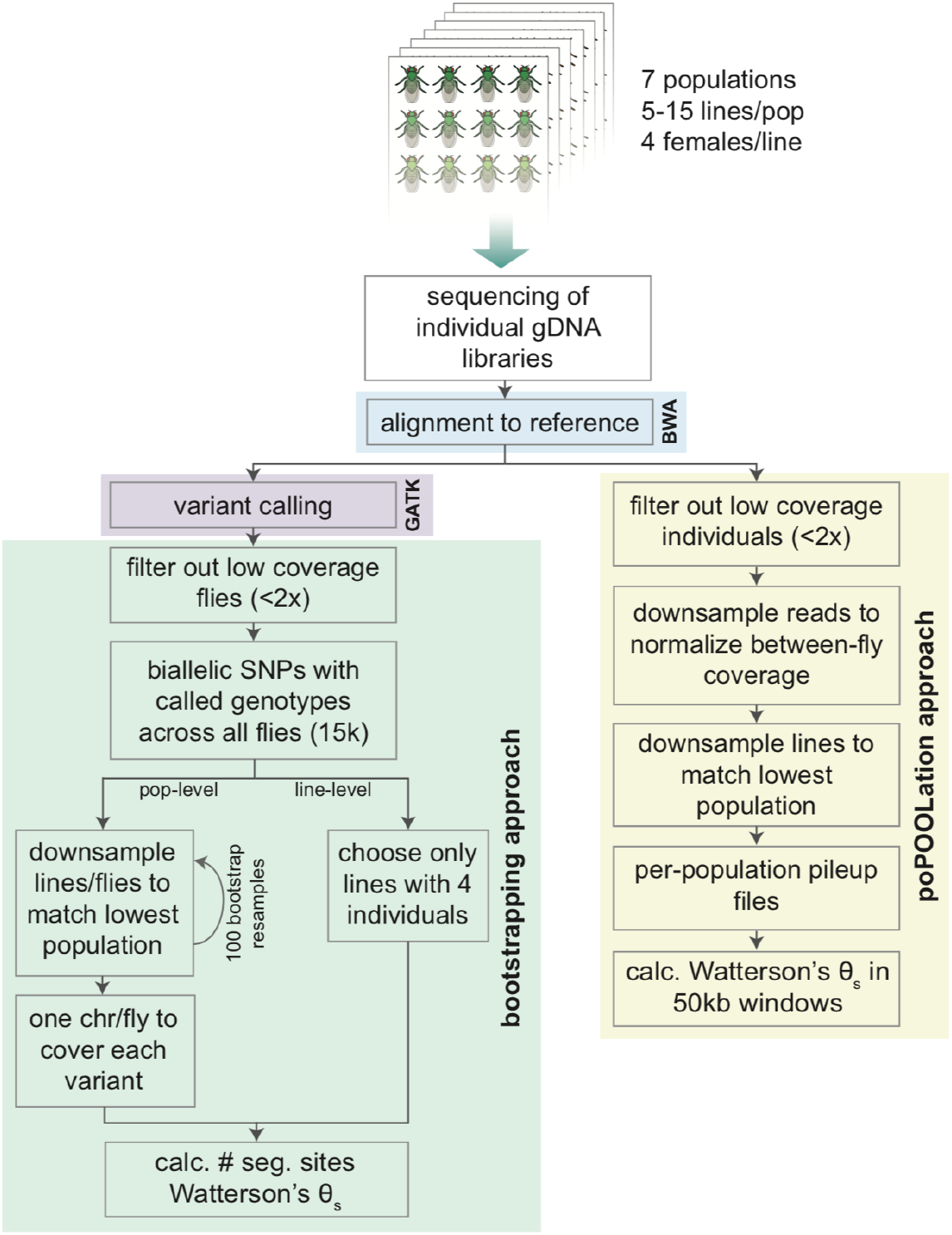
Flowchart of the two approaches used to estimate Watterson’s θ_s_.

## Supplementary discussion

Flies from FL exhibited patterns of thermal preference that were, depending on the experiment, both consistent and inconsistent with the model’s prediction that flies from this region would exhibit adaptive tracking. Specifically, these animals exhibited high heritability in thermal preference, consistent with an adaptive-tracking strategy (in bet-hedging animals, variation does not have a genetic basis) (Figs 2d, 4b). However, the dynamics of mean thermal preference in FL were more consistent with bet-hedging than adaptive tracking (Fig. 2d). The model’s prediction that adaptive tracking is favored in FL is rooted in that region’s year-round high temperatures, which produce (in the model) strong selection for colder thermal preference coupled with dramatic declines in population, as warm-preferring flies are selected against (Supp. Fig. 1). We collected fewer flies in FL, compared to MA and VA, and a period of declining sampling coincided with the hottest days of the year, as well as declining population in the model. This is consistent with high temperatures leading to strong selection against warm-preferring flies. It is possible that our outdoor traps captured a biased sample of the population, if cool-preferring flies were sheltering in cool microclimates, such as human residences. However, we did not observe increased warm-preferring behavior among the flies we did collect, as might be expected if cool-preferring flies were selectively avoiding the traps. An alternative hypothesis to explain the drop in collected flies is seasonal migration. Perhaps *D. melanogaster* retreats northward in FL summers, in a pattern mirroring seasonal repopulation of the northern limits of *Drosophila simulans*^1^.

When examining thermal preference heritability and variability, we were cognizant that genetic diversity could affect our estimates. We evaluated genome-wide levels of variation in individuals from isofemale lines by directly computing Watterson’s θ_s_ (with bootstrapping to estimate confidence intervals) as well as using the PoPoolation software package (Supp. Fig. 9). With neither method, did we see a strong relationship between θ_s_ and variability. We did observe a significant positive relationship with heritability (Supp. Fig. 3). This trend may be due to a geographic covariate. Both heritability and θ_s_ are higher in our south-eastern populations (TX and FL), in contrast with the northern populations. Higher θ_s_ in southern latitudes has been found in previous studies^2–4^. FL was an influential point in the relationship between θ_s_ and heritability, given that it was the only sampling site where we observed relatively high heritability of thermal preference. All in all, we are uncertain whether there is a causal relationship between θ_s_ and heritability, or if their correlation is imparted by geographic variation.

